# Spatial and temporal variation in farmland bird nesting ecology: Implications for effective Corn Bunting *Emberiza calandra* conservation

**DOI:** 10.1101/2024.09.02.610792

**Authors:** Nils Anthes, Julia Staggenborg, Markus Handschuh

## Abstract

Farmland bird populations continue to decline despite intense conservation efforts, possibly also because of incomplete knowledge on the drivers of local productivity. We therefore investigated spatial and temporal variation in nest site selection, breeding phenology, and nest survival for 225 nests of the Corn Bunting, a species of conservation concern in Central and Western Europe, in cropland-dominated, mixed, and grassland-dominated landscapes in SW Central Europe. Nesting phenology spread from April to August, started earlier at lower altitudes and progressed from grassland habitats to arable crops and eco-scheme flower fields. Most nests were placed in cultivated land, with substantial variation between landscape types but also between years within sites, so that large fractions of breeding attempts are prone to fail through land use operations in particular years. When nests were individually protected from land use, ‘apparent’ and ‘Mayfield’ survival rates differed substantially between nest habitats, with lowest survival in 2^nd^ year flower fields and highest survival in fallow grassland. Accounting for land use survival from patch-specific mowing, harvesting, and grazing dates, ‘total Mayfield nest survival’ estimates dropped by about half to 21 %, 13 %, and 20 % for (hay) meadows, alfalfa / clover-grass leys, and pastures, respectively, which held about 2/3 of nests in mixed landscapes. To enhance productivity beyond the thresholds required for local population persistence, we propose refined conservation schemes that improve survival within production farmland, best coupled with the development of prolific “Corn Bunting Landscapes”.

## 1 Introduction

Farmland bird populations continue to decline despite substantial conservation efforts (Burns et al. 2021, Rigal et al. 2023). Failed recovery may result from incomplete knowledge of the factors that limit population growth (Green 1995), such as large-scale determinants of distribution, abundance and population connectivity, the impact of land use or weather extremes on reproduction and survival, or local requirements for food or safe nesting sites (Crick et al. 1994). Since often at least some of that information is unavailable, conservation programmes tend to extrapolate from findings in other regions or species (Sutherland 2022), thus risking to implement locally insufficient or inappropriate measures.

The problem aggravates when conservation recommendations derive from correlative evidence that is mistaken for causation (Green 1995, Josefsson et al. 2020). For example, associations of territory placement with habitat composition can be spurious when populations in suboptimal habitats carry an extinction debt (Kuussaari et al. 2009, Hylander & Ehrlén 2013). They may also fail to identify nest habitats or other limitations to local productivity when driven by male song post availability (Van Horne 1983, Vickery et al. 1992). Habitat association studies therefore benefit from surveys of nest locations, hatching and fledging rates, and reproductive success (Green 1995, Sutherland 2022).

An example where incomplete information on breeding ecology may impede conservation is the Corn Bunting, *Emberiza calandra*. This farmland bird’s global population concentrates in Europe (Burfield et al. 2023) with ongoing declines in central and northwestern Europe (e.g., Crick et al. 1994, Anthes et al. 2017, Comolet-Tirman et al. 2021). Many Corn Bunting conservation schemes draw from observations in individual-rich or stable populations in vast open agricultural landscapes with > 10 % proportions of set-aside cropland, as in eastern Central Europe (Fischer 1999, Goławski & Dombrowski 2002, Schmidt et al. 2022). Given that territory-holding males associate with set-aside farmland also in western Central Europe (Meichtry-Stier et al. 2014, Burgess et al. 2015, Schmidt et al. 2022, Staggenborg et al. 2024), perennial fallows and eco-scheme flower fields are a key component of Corn Bunting conservation. However, this association considers neither the pairing status of territorial males, which is relevant given widespread polygyny in Corn Bunting (Hartley et al. 1993), nor nest locations, which may be clumped (Hegelbach 1997) and concentrate outside the ‘safe’ set-aside, nor breeding success, which may vary between different types of non-productive land.

It also remains unclear whether the positive effects of large field sizes and set-aside documented for eastern Central Europe can be mimicked in regions with more productive soils, higher crop diversity, smaller field sizes, and attractive alternative nesting habitats such as conventional cereals and clover-grass leys (Setchfield et al. 2012, Stein-Bachinger & Fuchs 2012) or meadows harvested amidst the breeding season (Suter et al. 2002, Broyer et al. 2014). For example, territory density and hatching success in French grassland populations only recovered with late mowing on ≥ 50 % of the local meadow area (Broyer et al. 2014, Broyer et al. 2016). In the central German Hainich National Park, a substantial part of the large and increasing Corn Bunting population inhabits permanent cattle and horse pastures with low stocking densities, where conflicts between nesting and land management are negligible given stable and predictable vegetation structures throughout the breeding season (Handschuh & Klamm 2022).

This study connects spatial and temporal variation of Corn Bunting nesting ecology with land use practices as a plausible driver for ongoing population declines. Covering several of the remaining marginal Corn Bunting populations in SW Central Europe, we first characterised variation in nest site selection along a gradient from cropland-to grassland-dominated landscapes. Second, we quantified Corn Bunting nesting phenology and its variation among regions, years, and habitats. Finally, we associated nest survival with nest habitat, season, and plot-specific mowing, harvesting, and grazing schedules. Based on these findings, we evaluate whether Corn Bunting conservation requires habitat- or landscape-specific adjustments to improve productivity in western Central Europe.

## 2 Methods

### 2.1 Study areas

Seven study areas in SW Germany and one in N Switzerland (Figure 1) comprised typical Corn Bunting nesting habitats spanning from landscapes dominated by extensive grasslands in the Rhine River floodplains to intensive arable farmland on loess plateaus; one study site (Rottenburg) comprised cattle pasture under rotational grazing with high stocking densities during the Corn bunting nesting season (site characterisations in supplemental Figure S1, Table S1). Most nest surveys took place between 2018 and 2020 (Table S1). In each study area and year, we mapped land use types (crop types and landscape elements) using QGIS (version 3.16). Between late April and mid-July, we recorded patch-specific land use dates for mowing, grazing, or (fodder) crop harvest; cereal harvest (of winter barley) rarely commenced before early July.

**Figure 1.**
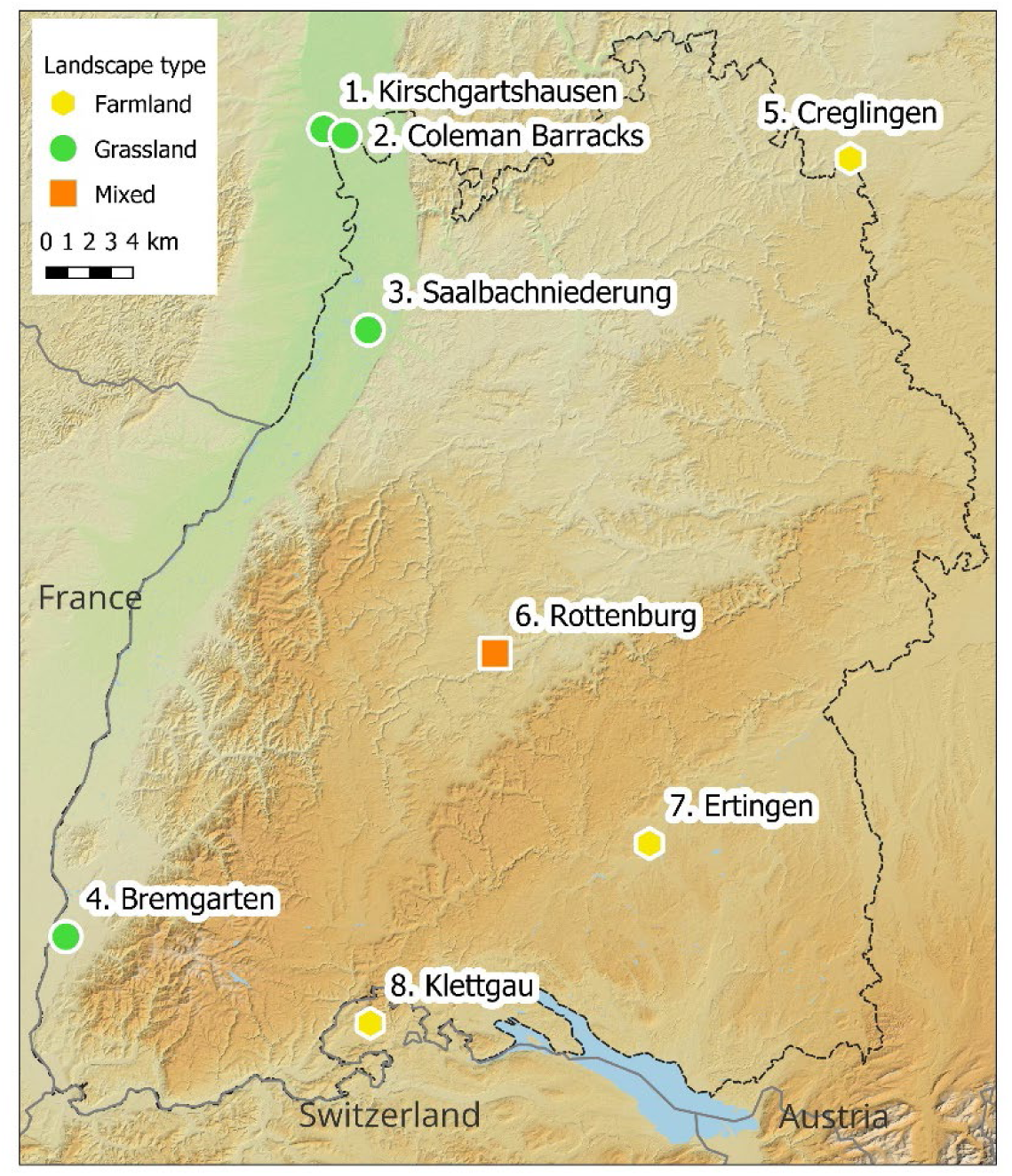
Location of the seven study areas. Labels link to supplemental Figure S1 and Table S1. Digital elevation model from European Union Copernicus data (EU-DEM layers).

### 2.2 Nest habitat and clutch parameters

We recorded the following parameters for 225 nests whose exact locations were derived from repeated observations of nest building activity at identical positions (n = 66 nests), repeated provision of nestling food to identical positions (n = 60 nests), or confirmation of nest sites through nest visits (n = 99 nests):

i. *Nest habitat:* Crop or land use category at the nest location (all nests).
ii. *Nest height*: Height of the upper nest rim above the ground (cm; visited nests),
iii. *Vegetation cover* and *Vegetation height*: Horizontal cover (5 % steps) and dominant height (10 cm steps) of the herbaceous vegetation estimated visually within 1 m of visited nests.

At visited nests, we further recorded the number of eggs and the number and age of nestlings in days (day 1 = hatching day). Nestling age was estimated according to a calibrated picture series from local nestlings of known age (supplemental Figure S2).

### 3.2 Nesting phenology

To compare nesting phenology with land use activities, we derived nest-specific timespans from nest building to full fledging from objective time points as available: nest building date, first egg date for incomplete clutches (backdated assuming a 1-day interval between successive eggs), and hatching date from nestlings of known age (available for n = 155 nests). We accepted less precise estimates when nests were visited only once during incubation, or when we confirmed nestling provisioning without nest visits. We assumed that such observations occurred midway in the incubation or nestling stage, respectively, and thus estimated their start 6 days before the observation, accepting a maximum estimation error of ± 6 days (n = 63 nests). From these dates, we extrapolated dates of first nest building, first egg laying, incubation, hatching, nest leaving, and full fledging based on literature data (Gliemann 1973, Boschert 1997, Hegelbach 1997) and own observations on the mean duration of each phase (supplemental Figure S3) rounded to the next full day to add a small buffer for delays that may occur in each nesting phase:

i. *Nest building*: 3 days (typically 2–4 days)
ii. *Egg laying*: 6 days (adjusted to full clutch size where known)
iii. *Incubation*: 13 days after clutch completion (typically 11–14 days)
iv. *Nestlings*: 12 days (9–13 days)
v. *Jumplings:* 12 days. Corn Bunting chicks leave the nest before they are able to fly (Hegelbach 1997). We term them ‘jumplings’ until fully fledged and capable of flight. In response to danger, jumplings typically remain stationary in the vegetation and may thus be killed by harvest or mowing given low cutting and high suction of modern land use machines. Previous studies imply that jumplings need at least 10–15 days after leaving the nest before switching from a ‘freeze’ to a flight escape response to approaching threats (Gliemann 1973, Perkins et al. 2013).

We analysed variation in nesting phenology with *first egg date* (FED) as the response variable and *study region* (four levels, supplemental Table S1) as a fixed predictor using generalized linear mixed effects models in the *glmmTMB* package (version 1.1.8, Brooks et al. 2017) in R (version 4.3.1, R Core Team 2023). Furthermore, we compared FED among *nest habitat* types (up to nine levels, see Figure 3) separately for two regions with sufficiently large sample sizes, Rottenburg and the Rhine River valley. These and all further models contained *study year* as random intercept to account for between-year variation in mean FED.

For model assessment according to Santon et al. (2023), we inspected residuals standardized for their distribution family regarding independence of fitted values and homogeneity across predictor variables and conducted posterior predictive checks on model-simulated data for dispersion, zero inflation, and distribution relative to observed data. FED model assessment favoured a Gaussian distribution, after excluding nests for which observations implied second or replacement broods because these generated extreme values that were insufficiently captured by our models.

We report coefficient and effect size estimates with their compatibility intervals, but refrain from presenting *P* values and their associated evaluation of binary null hypotheses in accordance with recommendations for unbiased statistical reporting (Halsey et al. 2015, Berner & Amrhein 2022).

### 3.3 Apparent nest survival and land use survival

We classified nests as ‘successful’ when adults were observed feeding at least one jumpling clearly outside but in the vicinity of a nest found active at least once between nest building and nestling stage.

*Apparent nest survival* quantifies the fraction of observed nests in which chicks reached the jumpling stage. This includes benefits of nest-specific contractual conservation agreements to postpone land use, i.e., any agricultural operation that removes potentially nest-holding vegetation through mowing, harvesting, or grazing. We therefore additionally quantified *land use survival* in the absence of nest protection as the fraction of study years in which a given nests breeding period did not overlap with observed, patch-specific land use dates, excluding years where contractual conservation agreements postponed land use on that patch.

*Apparent nest survival* and *land use survival* both overestimate true nest survival because early nest failures are underrepresented when nest detection is biased towards late incubation or the nestling stage (Mayfield 1975). Nevertheless, we took advantage of the maximum possible nest sample (supplemental Table S1) for comparative purposes, assuming similar nest detection rates (and thus similar bias in survival estimates) across nest habitats and regions.

Statistical analyses followed the linear modelling approach explained above, with *nest habitat* (9 levels) as the predictor variable and *study year* as random intercept. Model assessment favoured a logit-link binomial error distribution for *apparent nest survival* and a logit-link betabinomial error distribution for the mildly overdispersed *land use survival*.

### 3.4 Mayfield nest survival

Mayfield estimates of daily nest survival rates (DSR_nest_) remove the bias in apparent nest survival because they integrate the time each nest has been under survey (exposure duration, Mayfield 1975, Johnson 1979) and allow for variation in survival rates with nest age (Rotella et al. 2004, Weiser 2021). We analysed *daily nest survival rates* (DSR) for 93 nests visited at least twice during incubation or nestling phases, again including benefits of nest protection. We implemented Mayfield logistic regression in MARK (White & Burnham 1999), accessed through the R package RMark (Laake 2013), with nest outcome as the binary response variable (Dinsmore & Dinsmore 2007).

To minimize predictor collinearity, we conducted hierarchical modelling following Rotella et al. (2004) (supplemental Table S2). First, we screened for informative predictor variables among time and region predictors (set 1) and among nest and brood characteristics (set 2). Set 1 contained *Hatching Day* (days after earliest hatching observed in the dataset), *Time* (day of the year since our earliest nest recording), *Nest Age* (days after incubation start), study *Year*, and study *Region* (2 categories: Rottenburg and others). Set 2 contained *Nest Habitat* (simplified to seven levels to avoid small sample sizes: cereal fields combined with other crops, pasture combined with fallow grassland), *First Brood* (first versus replacement or second broods), *Clutch Size* (n eggs per clutch), *Nest Height*, *Vegetation Cover*, and *Vegetation Height*. Second, we combined the most informative predictor(s) per subset in a final model set, selecting predictors within 2 ΔAICc of the best performing model and retaining nesting habitat as our core predictor.

We derived mean DSR-estimates per nest habitat at median covariate levels and from these estimated nest survival across the entire incubation (13 days) and nestling (12 days) periods as DSR raised to the power of 25 (Johnson 1979).

## 3 Results

### 3.1 Nest habitat and clutch parameters

Nests were predominately placed in cultivated land (77 %), but habitats varied greatly among study regions (Figure 2a). In the cropland-dominated Creglingen region, most nests were in conventional winter wheat and barley. Mixed farmland landscapes had nests primarily in perennial eco-scheme flower fields in Ertingen/Klettgau, but showed a more even spread across meadows, alfalfa / clover-grass leys, cereals, and pastures in Rottenburg. Nests in the grassland-dominated Rhine River valley were mostly in fallow grassland and meadows. The large sample of nests from Rottenburg revealed between-year variation in nest habitats (Figure 2b), with a dominance of clover-grass versus cereal fields versus meadows alternating between successive years.

**Figure 2.**
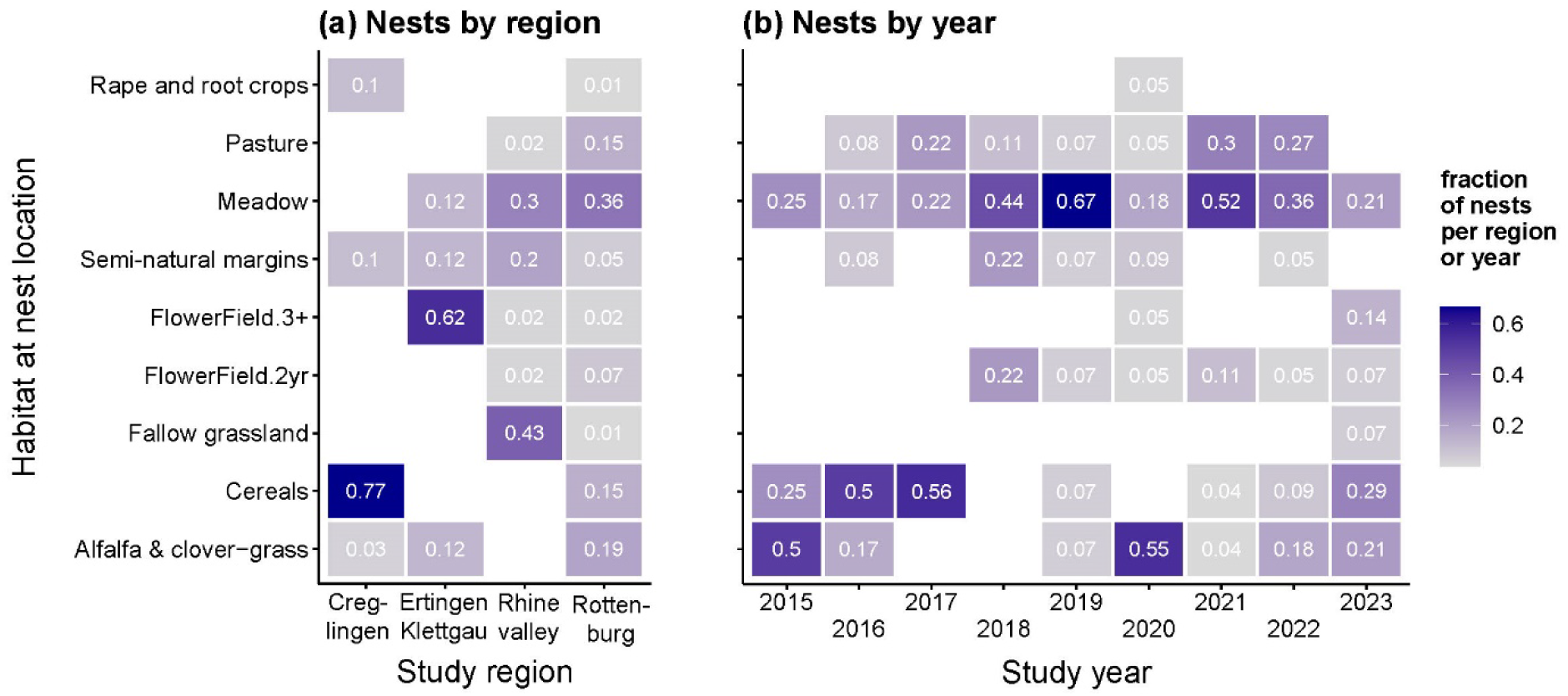
Corn Bunting nest habitats per study region (left panel) and per year at Rottenburg (right panel). Values show the proportion of nests per region or year. For sample sizes see supplemental Table S1.

Average clutch size was 4.49 eggs (95% CI: 4.08–4.95, n = 93 full clutches), without indications for variation with first egg date, study region, or nest habitat (Statistical Supplement A). Hatching rate, i.e., the fraction of eggs that hatched, was 0.88 on average (95% CI: 0.84–0.91, n = 70 clutches with hatching information). We found no variation in hatching rates with first egg date, clutch size, or study region, but hatching rates in eco-scheme flower fields tended to be lower than in other nesting habitats (Statistical Supplement A).

### 3.2 Nesting phenology

First egg dates (FED) spread between 22 April and 13 July and varied between study regions (Figure 3a). Earliest egg laying occurred in the Rhine valley lowlands (predicted mean FED: 20 May; 95% compatibility interval CI: 14–27 May), followed by Rottenburg (25 May, CI: 21–30 May), Ertingen/Klettgau (05 June, CI: 25 May–16 June) and Creglingen (07 June, CI: 31 May–13 June). Most pairwise differences were statistically robust (Statistical Supplement B).

**Figure 3.**
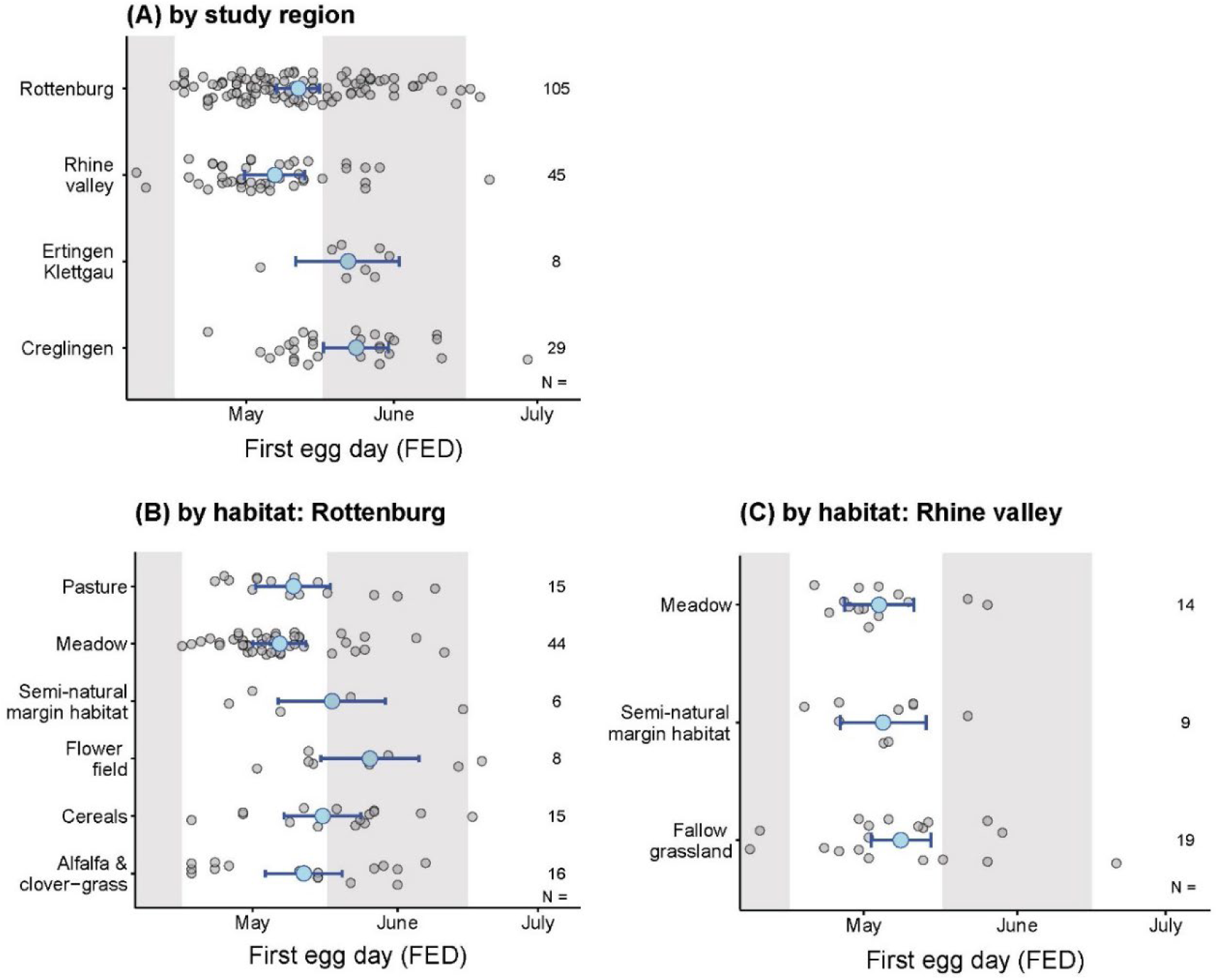
Variation in Corn Bunting first egg dates (FED) between regions (a) and nest habitats in Rottenburg (b) and the Rhine valley (c). Grey dots are raw data, large blue dots and flags are model-predicted mean values with 95 % compatibility interval. Estimated pairwise differences are given in Statistical Supplement B.

In Rottenburg, egg-laying was earliest in ‘grassland-like’ habitats (meadows and pastures), but some early FED also occurred in clover-grass leys (Figure 3b). Late egg laying occurred in cereal fields, semi-natural margin habitats, and perennial eco-scheme flower fields. Mean FED in flower fields was 19.3 (CI: 8.9–29.6) days later than in meadows (Statistical Supplement B). In the Rhine valley, egg laying dates were rather similar in all grassland nest habitats (Figure 3c, Statistical Supplement B).

### 3.3 Apparent nest survival and land use survival

Apparent nest survival varied substantially between nest habitats (Figure 4a, Table 1). It was highest in fallow grassland, followed by oilseed rape and root crops, and eco-scheme flower fields ≥ 3 years after sowing. Lowest apparent nest survival (< 60 %) occurred in flower fields in year 2 after sowing, and in alfalfa / clover-grass leys. These estimates contain nest protection measures (postponed mowing, harvesting, or grazing), which were initiated for 43 out of 74 nests in meadows, 25 out of 30 nests in alfalfa/clover-grass leys, and 19 out of 22 nests in pastures.

**Figure 4.**
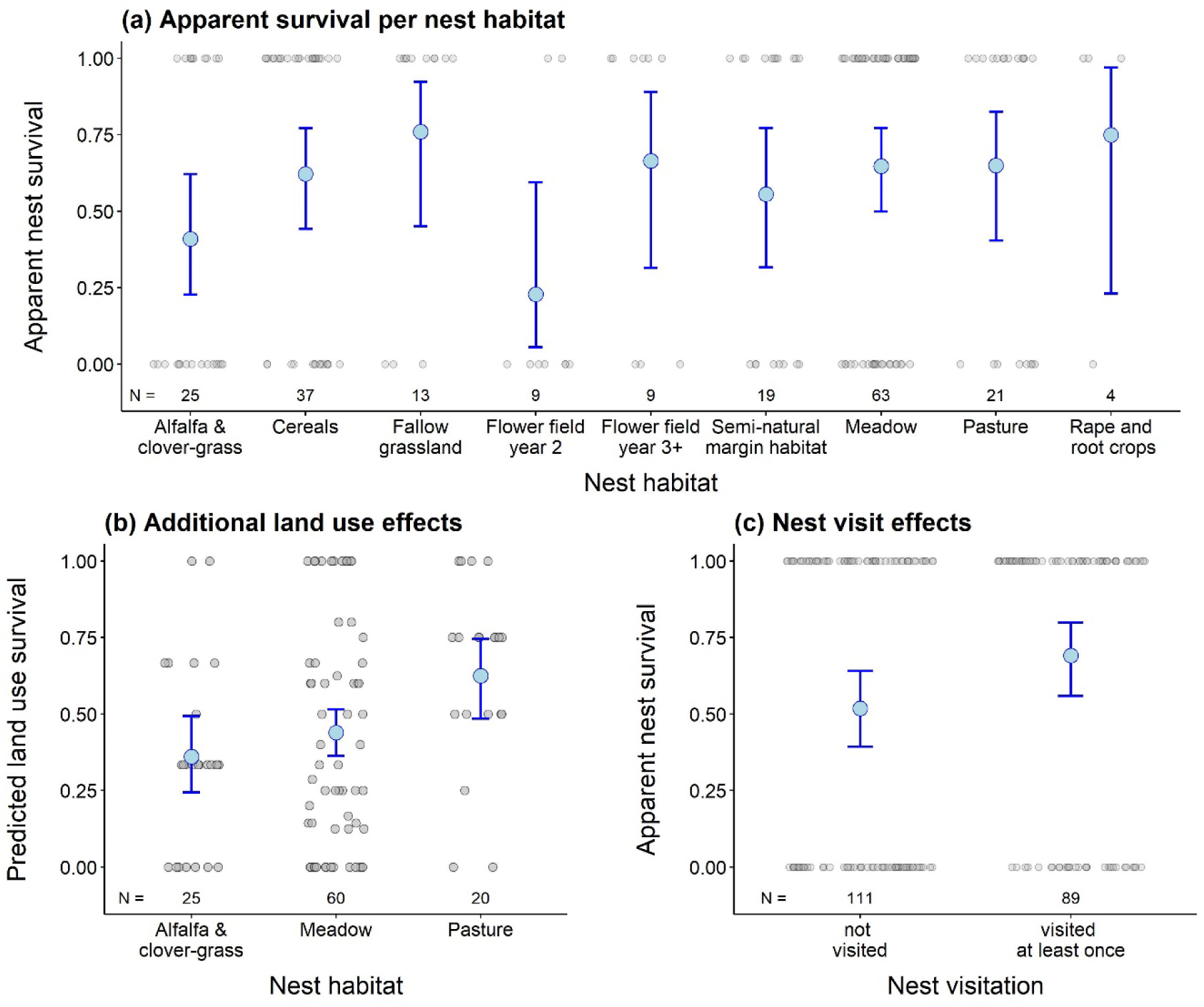
Variation in Corn Bunting nest survival. Panel (a) shows apparent nest survival per nest habitat, which includes survival benefits from contractual nest protection. Panel (b) shows land use survival, i.e., probabilities of completing the nesting period without collision with harvest, mowing, or grazing of the same patch (Figure 5). Panel (c) compares apparent nest survival between visited and unvisited nests. Grey dots display raw data, large blue dots and flags model-predicted means with their 95 % compatibility interval. Estimated pairwise differences are in Statistical Supplement C.

**Table 1.** Nest survival estimates per nest habitat. ‘Apparent survival’ as given in Figure 4a, ‘Land use survival’ as given in Figure 4b, Mayfield daily nest survival (DSR) as predicted for median hatching date per nest habitat. From the latter values we calculated Mayfield survival for the entire nesting period as DSR^25. Sample sizes are given in Figures 4, 6.

| Nest habitat | Apparent survival |  | Land use survival |  | Mayfield DSR |  | Mayfield survival |  |
| --- | --- | --- | --- | --- | --- | --- | --- | --- |
|  | mean | 95% CI | mean | 95% CI | mean | 95% CI | mean | 95% CI |
| Fallow grassland | 0.76 | 0.45-0.92 | - | - | 0.956 | 0.905-0.980 | 0.326 | 0.083-0.608 |
| Pasture | 0.65 | 0.40-0.83 | 0.62 | 0.49-0.75 | [Fallow grassland + pasture combined] |  |  |  |
| Flower field: year 2 | 0.23 | 0.06-0.59 | - | - | 0.906 | 0.787-0.962 | 0.085 | 0.003-0.379 |
| Flower field: year 3+ | 0.66 | 0.32-0.89 | - | - | 0.938 | 0.823-0.980 | 0.200 | 0.008-0.601 |
| Semi-natural margin habitats | 0.56 | 0.32-0.77 | - | - | 0.967 | 0.903-0.989 | 0.436 | 0.078-0.767 |
| Meadow | 0.65 | 0.50-0.77 | 0.44 | 0.36-0.52 | 0.972 | 0.946-0.985 | 0.486 | 0.251-0.688 |
| Alfalfa / clover-grass | 0.41 | 0.23-0.62 | 0.36 | 0.24-0.49 | 0.961 | 0.855-0.990 | 0.366 | 0.020-0.781 |
| Cereals | 0.62 | 0.44-0.77 | - | - | 0.955 | 0.870-0.986 | 0.319 | 0.031-0.696 |
| Rape, root crops | 0.75 | 0.23-0.97 | - | - | [Cereals, rape and root crops combined] |  |  |  |

*Land use survival,* i.e., the fraction of nests that would have their young fledged before land use on the same patch in the absence of nest protection (Fig. 5), was approx. 36 % for nests in alfalfa / clover-grass leys, 44 % in meadows, and 62 % in pastures (Figure 4b, Table 1). We cannot provide similar estimates for cereal fields because observed harvest dates almost never overlapped with active Corn Bunting broods, so cereal harvest was never postponed for nest protection. Thus, estimated apparent nest survival in cereal fields (approx. 62 %, Figure 4a, Table 1) includes potential harvesting effects.

**Figure 5.**
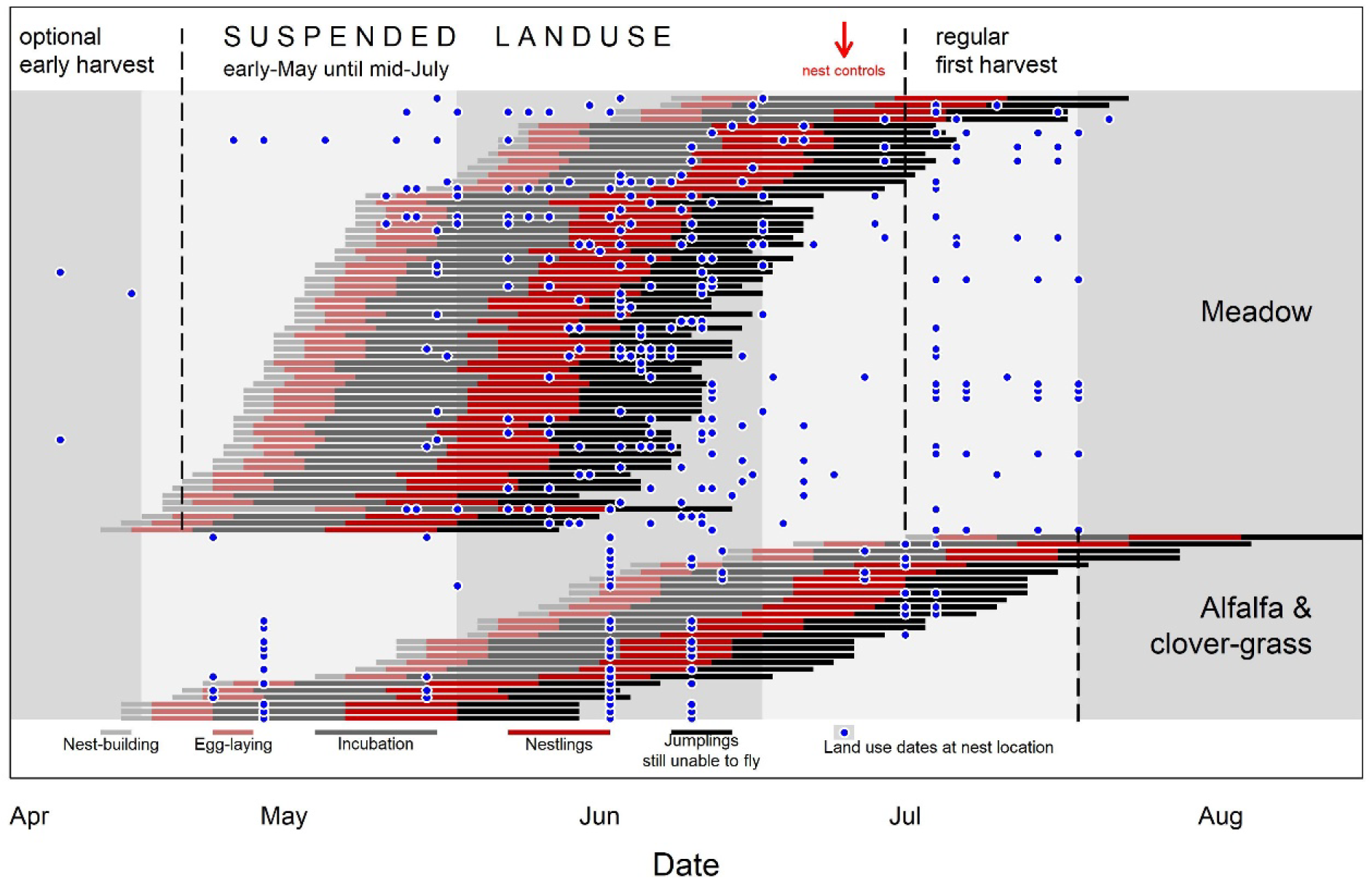
Nesting periods of individual Corn Bunting nests in meadows and alfalfa / clover-grass leys. Typical spans per breeding section are given in supplemental Figure S2. Blue dots indicate observed land use dates per individual field.

We found no indication that nest visits affected nest success (Figure 4c).

### 3.4 Mayfield nest survival

Mean daily nest survival rate (DSR) was 0.960 (95 % CI: 0.0.943–0.971), implying that nests had an average chance to survive the 25-day incubation and nestling period of 0.357 (CI: 0.220–0.485). DSR tended to decline with hatching date (Figure 6a) and nest height above ground (Figure 6b). Small sample sizes prevented a robust differentiation of DSR between nest habitats or regions (supplemental Table S2). Yet, best available estimates suggest low DSR for eco-scheme flower fields, particularly in their second year after sowing, and highest DSR in semi-natural margin habitats and meadows (Table 1, Figure 6c, d).

**Figure 6.**
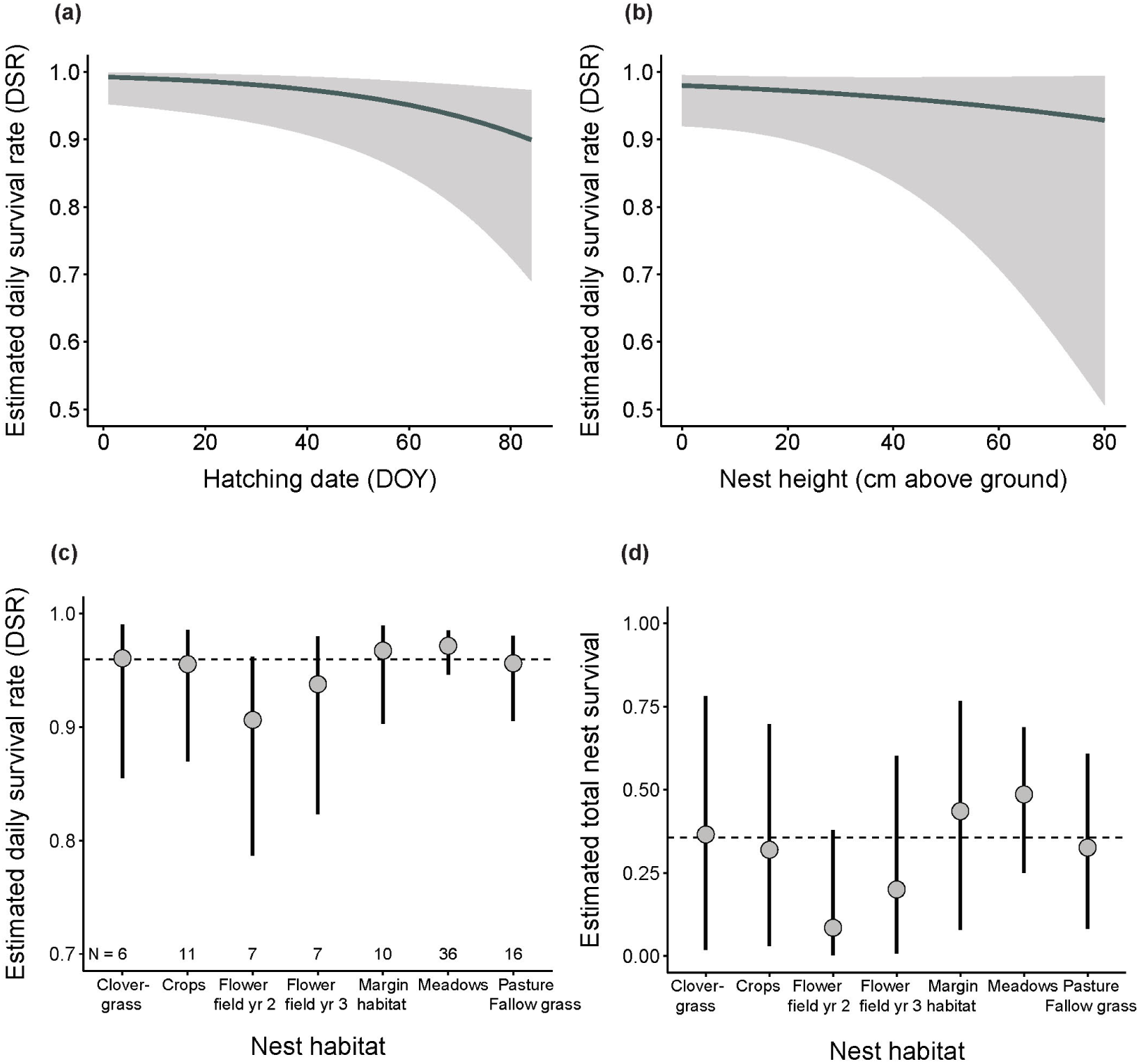
Mayfield estimates of daily nest survival rates (DSR) for hatching date (a; days since earliest recorded hatching), nest height (b), and nest habitat (c), and the estimated total survival during the 25-day nesting period per nest habitat (d) derived from (c). Dotted lines show overall mean survival rates in (c) and (d).

These Mayfield estimates still include nest protection benefits through postponed mowing, harvesting, or grazing. Multiplying them with land use survival estimated above (Tab. 1), our data suggest that just 13.2 % of nests in alfalfa / clover-grass leys, 21.4 % in meadows, and 20.2 % in pastures would have been successful, on average, in the absence of individual nest protection.

## 4 Discussion

Our study highlights several challenges for effective conservation of farmland birds with ground nests in cultivated land such as Corn Bunting.

First, nesting phenology spread widely between study sites and nest habitats. Earliest nests were built in April and last chicks fledged in late August. Nesting commenced earlier at lower altitudes and – within study sites – progressed from grassland habitats (meadows and pastures) to arable crops and eco-scheme flower fields. This sequence matches findings in British and French populations (Perkins et al. 2013, 2015, Setchfield et al. 2012, Brickle & Harper 2002) and likely arises from a preference for dense and medium-tall sward structures that provide optimal nest concealment (Perkins et al. 2015).

Second, we found strong variation in nest habitats between landscape types and study years. Cereals, fodder crops, and set-aside dominated in cropland-dominated landscapes and extensive or abandoned grass habitats in grassland-dominated landscapes, consistent with literature (supplemental Table S3). In mixed landscapes, we found a rather homogeneous spread of nests among several habitats. Conservation management thus requires solid knowledge of local nesting ecology before conservation measures can be copied from other regions.

Importantly, nest habitats also varied substantially between years: Most nests were placed in ‘low risk’ habitats such as cereal fields in some years, but in ‘high risk’ habitats in other years where the nest-holding vegetation is harvested during the Corn bunting breeding season. The latter required frequent interventions for nest protection through contractual land use postponement in alfalfa/clover-grass leys, meadows, and pasture. Our data imply that Mayfield nest survival rates of 36.6 %, 48.6 % and 32.6 % for these habitats in the presence of nest protection dropped to just 13.2 %, 21.4 %, and 20.2 %, respectively, under regular land use. These findings stress the effectiveness of nest protection, which roughly doubled nest success in three core nest habitats, especially in mixed landscapes (Fig. 2).

Our overall mean Mayfield DSR (0.960) falls within the range (0.894–0.979) reported across crop types and regions in Britain (Aebischer 1999, Brickle et al. 2000, Setchfield et al. 2012, Perkins et al. 2013). However, our habitat-specific nest survival estimates for cereals and fallow grassland (0.32) are clearly below those reported for cereals and rough grass in Scotland (0.45–0.5, Perkins et al. 2013). Similar to findings from Scotland (Perkins et al. 2013), we found that DSR declined moderately as season progressed. Late broods more likely collide with mowing, grazing, or harvest, but these have minor impact in our dataset given protection of almost all known nests. We inferred plausible causes for 24 out of the 30 nest failures observed among 90 visited nests, with all except the first category dominating in late season nests: predation (12 cases), association with severe rain, cold or drought (7 cases) as recently increasing during central European summers (Hänsel et al. 2022), land use (3 cases of failed conservation intervention), tall vegetation that fell on the nest (1 case), and nestling starvation (1 case), possibly driven by a lack of insect food availability (Grames et al. 2023) during drought. DSR also tended to decline for above-ground nests, which in our study were restricted to tall vegetation without dense ground swards, such as young eco-scheme flower fields. While elevated nests tended to tip over, several nests also showed signs of predation by mammals, which can easily access and patrol such habitats.

We found comparably large clutches with solid hatching success (Hartley & Shepherd 1994, supplemental Table S3) and a mean size of successful broods (3.85 jumplings, n = 60 visited successful nests) exceeding values reported from Scotland (2.9, Perkins et al. 2013). We therefore argue that poor population recovery in our study areas is driven less by hatching success and brood size than by nest failures (as shown above), a low fraction of females that raise a second brood (Siriwardena et al. 2000, Brickle & Harper 2002, Setchfield et al. 2012), and likely habitat loss. Indeed, the average number of successful broods per male territory in our core study site ‘Rottenburg’ – a crude indicator of productivity – span from 0.30 to 0.71 in 2014–2023 (unpubl. data), never crossing the minimum value of 0.75 that would be required for population persistence assuming mean adult survival of 0.58, mean juvenile recruitment of 0.29 (both values from Perkins et al. 2013) and our mean output of 3.85 jumplings per successful nest.

### 4.1 Conservation and management implications

Meadows in grassland-dominated and mixed landscapes attracted a substantial fraction of nesting attempts but associated with a high failure risk from mowing, even in late-mown hay meadows. Perkins et al. (2013) also report low nest survival rates in meadows under conventional mowing (DSR 0.89) compared to delayed mowing (DSR 0.96). The difference was largely caused by mowing-induced nest failures of 66 %, closely matching our estimated ‘land use failure’ rate for meadows of 56 %. Delayed mowing can restore population productivity when implemented on 50–75 % of the meadow habitat that is relevant for the local Corn Bunting population (Perkins et al. 2013, Broyer et al. 2016). Fractions < 20 % only suffice when detailed prior knowledge allows targeting meadow patches with proven highest quality for, and thus reliable clustering of, Corn Bunting nests (Perkins et al. 2013). Non-targeted delayed mowing on such small land fractions likely fails because birds cannot distinguish early from late mown grassland at the onset of nesting and thus do not focus nest placement onto protected patches (Broyer et al. 2012).

Safe mowing dates depend on local nesting schedules, with 15 July and 01 August plausible options in regions with regular (Rottenburg, Figure 2) or late breeding (Scotland, S England, Brickle & Harper 2002, Perkins et al. 2013), respectively. While late mowing can allow the completion of late first broods, early replacements, and second broods, it may locally compromise yield or meadow habitat quality, potentially conflicting with regulations to protect Lowland hay meadows (habitat type 6510) under the EU-habitats directive. Conservation contracts should thus encourage early grazing or mowing before the onset of the local farmland bird breeding season for weed control (e.g., autumn crocus *Colchicum autumnale*), grass suppression, maintenance of yield, and prevention of tall grass swards that tend to collapse from wind exposure or rain. Such a regime partially mimics traditional land use where winter or spring pasture preceded the meadow period (Kapfer 2010), also to the benefit of insect diversity and abundance (Humbert et al. 2012, Bruppacher et al. 2016). We recommend late mowing as a complementary measure to landscape-scale restoration of safe nest habitats that are more attractive for Corn Buntings than meadows. Prime options include long-term set-aside (Staggenborg & Anthes 2022) and the transformation of meadows into permanent or seasonal (but not rotational) cattle or horse pastures with low stocking densities as a favourable Corn Bunting habitat (Handschuh & Klamm 2022).

Our findings confirm that alfalfa / clover-grass leys can act as an ecological trap since they attract nesting Corn Buntings and other farmland birds (Stein-Bachinger & Fuchs 2012, González del Portillo et al. 2022) but are associated with high risks of nest failure given harvest during the breeding season. Clover-grass is a key element of organic crop rotation for nitrogen fixation, weed suppression, fresh or silage livestock fodder, and bioenergy, typically covering 20-30 % of organic farmland. Its relevance will increase given EU aims to expand organic farming to 25 % by 2030 (Fetting 2020). We suggest contractual Corn Bunting conservation to ban clover-grass harvest between at least 01 May and earliest mowing dates proposed above, or to counsel farmers to integrate clover-grass leys into set aside requirements under EU regulations for good agricultural and environmental conditions (GAEC). Alternatives may include bi-annual no-harvest strips within regularly used clover-grass leys and cereal-legume mixes, with harvest not until August; this intervention, however, is currently untested and potential effects on Corn Bunting breeding success, nitrogen fixation and weed control remain to be established.

Finally, we found that early successional eco-scheme flower fields (year 2 after sowing) attracted only few Corn Buntings nests with then low success rates. Tall vegetation coupled with low sward density near the ground can increase nest predation rates (Perkins et al. 2015). In contrast, more mature flower fields – three or more years since sowing – not only represent preferred territory centres (Zollinger et al. 2013) and all-year foraging habitat for Corn Buntings (Rieger et al. 2022), but also provide safe nesting structures in patches with dense vegetation at ground level. Thus, depending on the chosen seed mix and the locally developing vegetation structure, late successional flower fields can qualify as a targeted Corn Bunting conservation measure (Staggenborg & Anthes 2022).

### 4.2 Conclusion

Our study highlights that effective farmland bird conservation requires solid knowledge of local nesting ecology and that conservation measures cannot necessarily be transferred between habitats, landscape types, or regions. Its long breeding season and variable nest habitat selection between regions and years challenge Corn Bunting conservation in intensive agricultural landscapes. We derive two key recommendations from our findings.

First, given a small fraction of nests in set-aside or non-productive fields (supplemental Table S4), effective Corn Bunting conservation must stabilize and improve nest survival on cultivated land. We suggest refined conservation schemes for alfalfa/clover-grass leys, meadows, and pastures to enhance local Corn bunting productivity. Future studies should evaluate the success of the proposed measures and elucidate the currently understudied collision of the Corn Bunting nesting period with cereal harvest and with mechanical weed control in organic crops.

Second, non- or low-productive areas (in a land sparing sense, Grass et al. 2021) may only boost Corn Bunting productivity when becoming a dominant feature in the landscape, e.g., in landscapes with extensive permanent pasture or substantial fractions of abandoned land (Handschuh & Klamm 2022). We suggest the development of such model “Corn Bunting Landscapes”, ideally associated with agricultural rewilding (Corson et al. 2022), where large fractions of land provide attractive nesting and feeding habitat for farmland birds.

A two-tier approach that increases the reproductive output of farmland birds within productive farmland while providing strategically placed species strongholds with prolific source populations may constitute the most promising approach to a recovery of farmland bird populations in line with the novel EU Nature Restoration Law.

## Acknowledgements

This research received administrative support through the regional councils of Freiburg, Karlsruhe and Stuttgart as well as local district administrations. Thorsten Teichert and ‘Verein Vielfalt e. V. Tübingen’ were indispensable in coordinating nest protection measures at our core study site Rottenburg. We further thank Clarissa Anders, Martin Boschert, Santiago Cruzes Vallo, Sabine Geissler-Strobel, Heiner Götz, Valentin Grom, Annika Hammerschmidt, Alexandra Ickes, Markus Jenny, Ariane Koch, Mathias Kramer, Franziska Lehle, Miriam Plappert, Timo Reischmann, Benjamin Reichelt, Mirjam Schöckle and Caroline Schuck for general support or contributions to field work.

## 5 Author contributions

Study conception (NA, MH), Data collection (MH, NA, JS), Data analysis lead (NA), Data analysis support (MH, JS), Manuscript lead (NA), Manuscript support (MH, JS).

## 6 Declarations

### Permits

Nest surveys were conducted under ringing permit to N.A. (ringer ID 0674 at Vogelwarte Radolfzell) issued within conservation permit Az. 55-8841.03; 8853.17 by Regierungspräsidium Karlsruhe (25.4.2016) and Corn Bunting conservation mandates of the Federal State Agencies of Tuebingen, Stuttgart, Karlsruhe, and Freiburg.

### Data availability

Raw data for all reported analysis are available via Zenodo (https://doi.org/10.5281/zenodo.12799271).

