## Supplemental Material for "Spatial and temporal variation in farmland bird nesting ecology: Implications for effective Corn Bunting *Emberiza calandra* conservation"

Preprint - version July 2024

### Supplemental Figures

#### Figure S1 – Habitat composition per study area

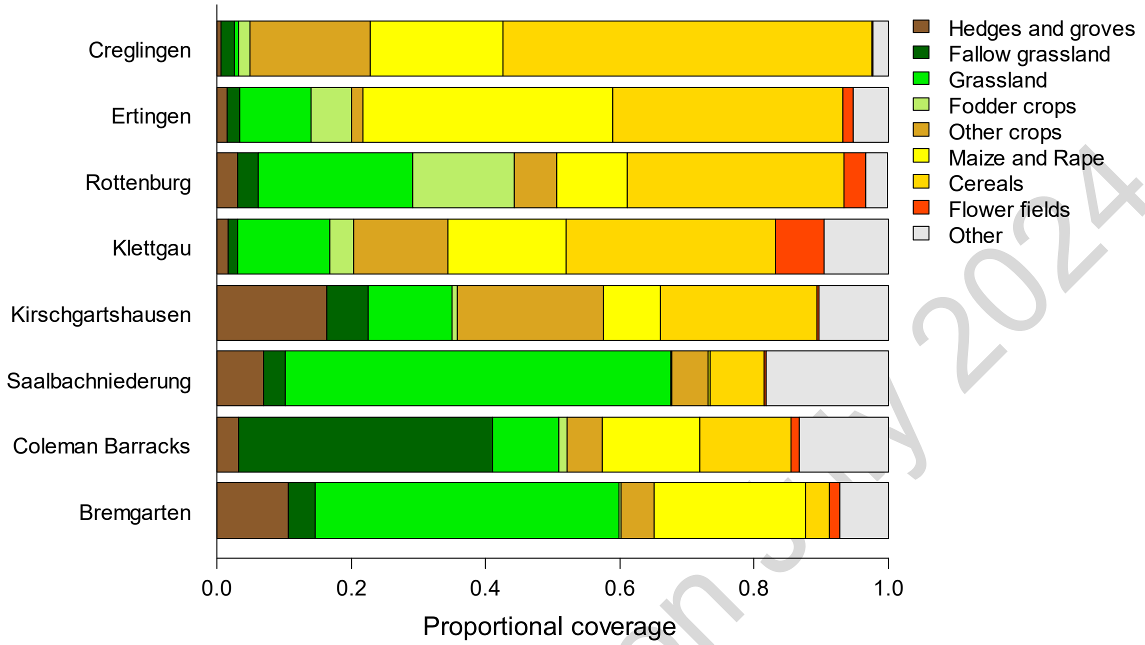

**Figure S1**. Habitat composition of the study areas by main land use types. Study areas are arranged from grassland-dominated (bottom) to cropland-dominated landscapes (top).

Preprtint - version July 2024

#### Figure S2 – Corn Bunting nestling age

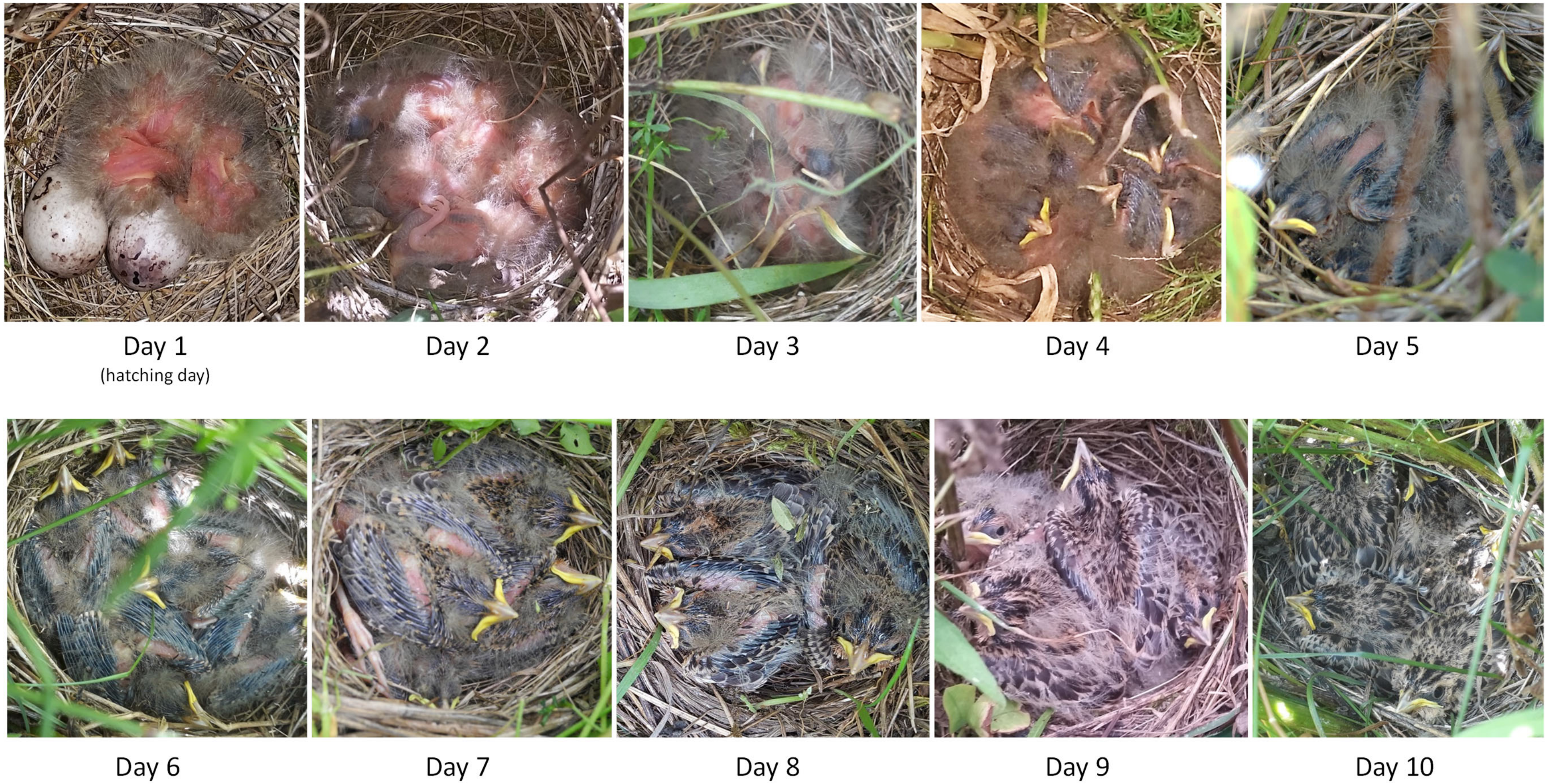

**Figure S2**. Calibrated pictures of Corn Bunting nestlings of known age post-hatching. Image © Nils Anthes & Markus Handschuh.

#### Figure S3 – Corn Bunting breeding phases

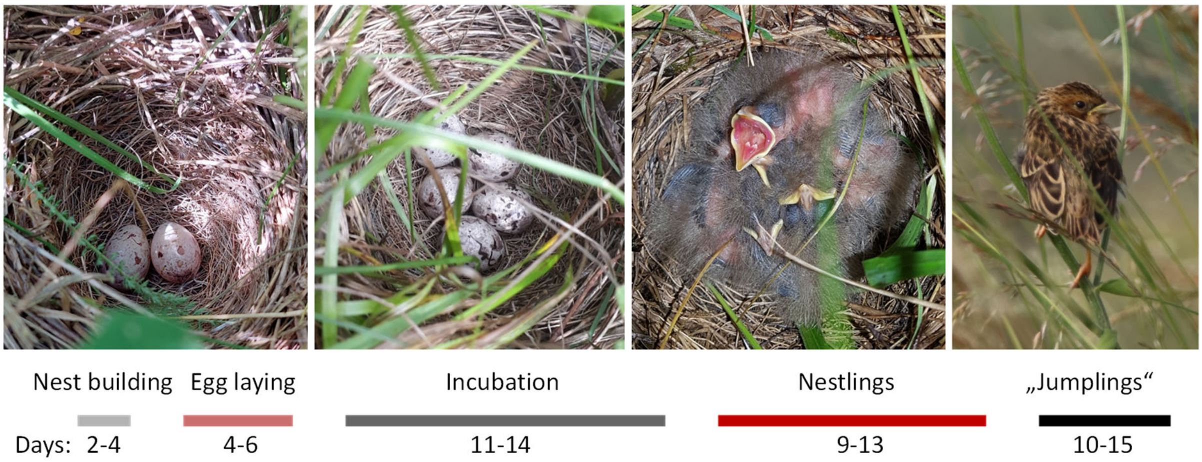

**Figure S3**. Corn Bunting breeding phases and their estimated duration. Image © Nils Anthes & Markus Handschuh.

Preprtint - version July 2

### Supplemental Tables

#### Table S1 – Study area: characteristics and sample sizes

**Table S1**. Characterisation of study areas and the number of nests per site that were integrated into the analysis of nesting habitats ('habitat'), nesting phenology ('phenology'), apparent nest survival ('app. surv.'), and Mayfield daily nest survival ('DSR'). Study areas with shared background colour were combined by 'Region' for statistical analysis.

| **Study area** | **Landscape type** | **Region** | **Area (ha)** | **Study years** | **Sample size (# nests)** | | | |
|---|---|---|---|---|---|---|---|---|
| | | | | | *habitat* | *phenology* | *app. surv.* | *DSR* |
| (1) Kirschgarts-hausen | Restored floodplain grasslands, in a forest and arable land matrix | Rhine valley grasslands | 387 | 2018-2020 | 12 | 12 | 10 | 2 |
| (2) Coleman Barracks | Extensive grasslands on airstrip, surrounded by arable fields | Rhine valley grasslands | 316 | 2018-2020 | 16 | 16 | 11 | 2 |
| (3) Saalbach-niederung | Mesotrophic meadows in arable landscape matrix | Rhine valley grasslands | 385 | 2018-2020 | 5 | 5 | 5 | 1 |
| (4) Bremgarten | Extensive grasslands on airstrip fields and adjacent arable fields | Rhine valley grasslands | 765 | 2018-2020 | 13 | 13 | 10 | 3 |
| (5) Creglingen | Intense arable land | Creglingen farmland | 650 | 2019-2022 | 30 | 30 | 23 | 8 |
| (6) Rottenburg | Extensive arable land, high proportions of organic farming, meadows and cattle pastures | Rottenburg mixed | 590 | 2014-2022 | 141 | 134 | 133 | 72 |
| (7) Ertingen | Intense arable land, dedicated farmland eco schemes concentrate in centre | Ertingen / Klettgau farmland | 439 | 2019 | 2 | 2 | 2 | 1 |
| (8) Klettgau | Intense arable land, dedicated farmland eco schemes concentrate in centre | Ertingen / Klettgau farmland | 305 | 2018-2019 | 6 | 6 | 6 | 4 |
| **Total** | | | | | **225** | **218** | **200** | **93** |

Preprint - ve

#### Table S2 – Model selection for Mayfield daily nest survival rates (DSR)

**Table S2**. Overview of MARK-models to predict variation in Daily nest survival rates (DSR). Model sets (a) and (b) served to extract predictor combinations for the combined model set (c). Within each model set, predictor combinations are sorted by AICc values. 'npar' is the number of estimated model parameters.

| Model set and predictors | npar | AICc | Δ AICc | weight | Deviance |
|---|---|---|---|---|---|
| **(a) Time and Region** | | | | | |
| HatchingDay | 2 | 144.68 | 0.00 | 0.41 | 140.66 |
| Time | 2 | 146.12 | 1.44 | 0.20 | 142.10 |
| HatchingDay + Time | 3 | 146.64 | 1.96 | 0.15 | 140.60 |
| HatchingDay + NestAge | 3 | 146.65 | 1.97 | 0.15 | 140.61 |
| HatchingDay + Time + NestAge | 4 | 148.59 | 3.91 | 0.06 | 140.53 |
| NestAge | 2 | 151.99 | 7.32 | 0.01 | 147.97 |
| (Intercept) | 1 | 153.30 | 8.63 | 0.01 | 151.30 |
| Region | 2 | 155.31 | 10.64 | 0.00 | 151.29 |
| Year | 7 | 161.28 | 16.60 | 0.00 | 147.11 |
| **(b) Nest and brood characteristics** | | | | | |
| FirstBrood | 2 | 149.71 | 0.00 | 0.48 | 145.69 |
| NestHeight | 2 | 150.56 | 0.85 | 0.32 | 146.54 |
| (Intercept) | 1 | 153.30 | 3.60 | 0.08 | 151.30 |
| Clutch Size | 2 | 154.11 | 4.40 | 0.05 | 150.09 |
| VegetationCover | 2 | 155.13 | 5.42 | 0.03 | 151.11 |
| VegetationHeight | 2 | 155.21 | 5.50 | 0.03 | 151.19 |
| NestHabitat | 7 | 160.71 | 11.00 | 0.00 | 146.54 |
| **(c) Combined models** | | | | | |
| HatchingDay | 2 | 144.68 | 0.00 | 0.29 | 140.66 |
| HatchingDay + NestHeight | 3 | 145.49 | 0.81 | 0.19 | 139.45 |
| HatchingDay + FirstBrood | 3 | 146.01 | 1.34 | 0.15 | 139.98 |
| HatchingDay + Time | 3 | 146.64 | 1.96 | 0.11 | 140.60 |
| HatchingDay + NestHeight + FirstBrood | 4 | 146.77 | 2.10 | 0.10 | 138.71 |
| HatchingDay + Time + NestHeight | 4 | 147.49 | 2.82 | 0.07 | 139.43 |
| HatchingDay + Time + FirstBrood | 4 | 148.02 | 3.34 | 0.05 | 139.96 |
| FirstBrood | 2 | 149.71 | 5.03 | 0.02 | 145.69 |
| NestHeight | 2 | 150.56 | 5.88 | 0.02 | 146.54 |
| (Intercept) | 1 | 153.30 | 8.63 | 0.00 | 151.30 |
| HatchingDay + NestHabitat | 8 | 153.70 | 9.02 | 0.00 | 137.48 |
| HatchingDate + NestHabitat + NestHeight | 9 | 154.84 | 10.16 | 0.00 | 136.57 |
| HatchingDate + NestHabitat + NestHeight + FirstBrood | 10 | 156.28 | 11.60 | 0.00 | 135.95 |
| HatchingDate + NestHabitat + NestHeight + Time | 10 | 156.89 | 12.21 | 0.00 | 136.56 |
| NestHeight + NestHabitat | 8 | 159.87 | 15.20 | 0.00 | 143.65 |
| NestHabitat | 7 | 160.71 | 16.03 | 0.00 | 146.54 |

Preprint - version July 2024

#### Table S3 – Literature overview: Corn Bunting nest habitats and nest parameters

**Table S3**. Summary of literature values on the distribution of Corn Bunting nesting habitats across regions and landscape types. Studies varied in the nomenclature and clustering of nest habitats. We allocated values to nest habitat categories as precisely as possible, and comment on specific cases where this was only partially possible.

| Source | LandscapeType | N nests | Winter cereals | Spring cereals | Fodder crops[1] | Root crops | Fallow & set aside[2] | Meadow intensive | Grassland extensive | Pasture | Margins[3] | Ruderal land | Bushland | Other | Clutch size | Hatch rate | Jump/succ | Jump/nest | Country | Region | Comment |
|---|---|---|---|---|---|---|---|---|---|---|---|---|---|---|---|---|---|---|---|---|---|
| Sacher & Bauschmann 2011 | cropland-dominated | **16** | 0.88 | 0.00 | 0.06 | 0.00 | 0.06 | 0.00 | 0.00 | 0.00 | 0.00 | 0.00 | 0.00 | 0.00 | - | - | - | - | Germany | central and S Hessen | 13 wheat, 1 barley, 1 alfalfa |
| Fehn 2021 | cropland-dominated | **40** | 0.53 | 0.00 | 0.20 | 0.03 | 0.18 | 0.00 | 0.00 | 0.00 | 0.00 | 0.00 | 0.00 | 0.00 | - | - | - | - | Germany | Zülpicher Börde region | |
| Hölker & Klähr 2004 | cropland-dominated | **27** | 0.56 | 0.04 | 0.07 | 0.11 | 0.15 | 0.00 | 0.00 | 0.00 | 0.04 | 0.00 | 0.04 | 0.00 | 4.3 | - | - | 2.75 | Germany | Hellwegbörde (NRW) | |
| Perkins et al. 2015 | cropland-dominated | **580** | 0.12 | 0.50 | 0.22 | (**) | 0.06 | | | (**) | 0.06 | (*) | 0.00 | 0.05 | - | - | - | - | Scotland | 4 regions in E Scotland | 'fodder crops`= forage grass. 66 in "other grass types such as non-rotational set-aside, field margins, newly sown grass", here split 1:1 between fallow and margins. 'Other' = Rape, Root crops, and Pasture (**). |
| Brickle & Harper 2002 | cropland-dominated | **120** | 0.33 | 0.36 | 0.00 | 0.00 | 0.15 | 0.00 | 0.08 | 0.00 | 0.08 | 0.00 | 0.00 | 0.00 | - | - | - | - | England | West Sussex | |
| Stein-Bachinger et al. 2010 | cropland-dominated (mostly organic) | **77** | 0.16 | 0.08 | 0.47 | | | | | | | | | 0.30 | 4.8 | 0.95 | - | 1.84 | Germany | Brandenburg | No further differentation by crop types |
| Gyllin 1965 (from Gliemann 1973) | cropland-dominated | **59** | 0.00 | 0.20 | 0.69 | 0.00 | 0.00 | 0.00 | 0.00 | 0.00 | 0.10 | 0.00 | 0.00 | 0.00 | - | - | - | - | Sweden | | |
| Eislöffel 1997 | cropland-dominated | **51** | | | | | 0.59 | | | | 0.10 | | | 0.31 | - | - | - | - | Germany | Rheinland-Pfalz | 'Other' not further differentiated |
| Setchfield et al. 2012 | cropland-dominated | **200** | 0.00 | 0.59 | 0.08 | (*) | 0.04 | 0.08 | 0.04 | 0.04 | 0.04 | | (*) | 0.10 | - | - | 2.78 - 3.33 | - | England | Cornwall | All Cereals = spring barley (often 'extensive'). 'silage & haylage': split 1:1 among fodder crops and intense meadows. 'extensive grassland' split 1:1 among abandoned pasture, margins, extensive coastal grassland, and set-aside. 'Others' contain root crops, bushes (*) and others. |
| Fischer 1999 | mixed | **106** | 0.04 | 0.01 | 0.06 | 0.00 | 0.49 | 0.06 | | | | 0.31 | | 0.04 | 4.7 | - | - | - | Germany | Brandenburg | |
| Suter et al. 2002 | mixed | **35** | 0.26 | 0.00 | 0.00 | 0.40 | 0.03 | 0.26 | 0.06 | 0.00 | 0.00 | 0.00 | 0.00 | 0.00 | - | - | - | - | Switzerland | Großes Moos | |
| Gliemann 1973 | mixed | **39** | 0.10 | | 0.05 | 0.00 | 0.31 | 0.36 | | 0.00 | 0.00 | (*) | 0.18 | 0.00 | 4.7 | - | 4.22 | - | Germany | Oberlausitz region | Bushland = Broom *Genista*, Black- and Raspberry *Rubus*, 'fallow' also contains ruderal land (*). |
| Boschert 1997 | various | **60** | 0.03 | 0.02 | 0.03 | 0.00 | 0.02 | 0.47 | | 0.00 | 0.32 | 0.12 | 0.00 | 0.00 | 4.5 | - | - | - | Germany | Baden-Württemberg | 'Ruderal land' contains nests in quarries and 'wasteland'. |
| Mildenberger 1984 | various | **129** | 0.37 | | 0.08 | 0.00 | | 0.52 | | | 0.00 | 0.03 | 0.00 | 0.00 | - | - | - | - | Germany | Rheinland region | |
| Hegelbach 1984 | grassland-dominated | **113** | 0.00 | 0.00 | 0.00 | 0.00 | 0.00 | 0.00 | 1.00 | 0.00 | 0.00 | 0.00 | 0.00 | 0.00 | 4.1 | | 3.20 | | Switzerland | Reußtal | 112 nests in extensive litter meadows, 1 in conventional meadow. |
| Ryves 1934 (from Gliemann 1973) | Coastal Heathland | **54** | 0.09 | | 0.00 | 0.00 | 0.00 | 0.00 | 0.89 | 0.00 | 0.00 | 0.02 | 0.00 | 0.00 | 3.9 | - | - | - | England | N Cornwall | study focused on coastal heathland, with most nests in heather *Calluna* and Common Gorse *Ulex europaeus*. |

[1] Fodder crops: mostly alfalfa, clovergrass, field grass, peas
[2] Fallows: temporarily unused cropland, also often termed 'set-aside'. Includes eco-scheme flower fields.
[3] Margins: Ditches, field margins, grasspaths, grass seams

**Sources for Table S3**

Boschert, M. (1997). Grauammer (*Miliaria calandra*). In Hölzinger, J. (eds), Die Vögel Baden-Württembergs. Band 3.2 - Singvögel 2, (pp. 831-860). Ulmer.

Brickle, N.W. and Harper, D.G. (2002). Agricultural intensification and the timing of breeding of Corn Buntings *Miliaria calandra*. Bird Study 49, 219-228.

Eislöffel, F. (1997). The corn bunting *Miliaria calandra* in south-west Germany: population decline and habitat requirements. In Donald, P.F. and Aebischer, N.J. (eds), The ecology and conservation of corn buntings *Miliaria calandra*. , (pp. 170-173). Joint Nature Conservation Committee, Peterborough.

Fehn, M. (2021). Brutökologie, Raumnutzung und Habitatwahl der Grauammer (*Emberiza calandra*) in der Zülpicher Börde NRW. MSc-Arbeit an der Universität Bonn.

Fischer, S. (1999). Abhängigkeit der Siedlungsdichte und des Bruterfolgs der Grauammer (*Miliaria calandra*) von der agrarischen Landnutzung: Ist das Nahrungsangebot ein Schlüsselfaktor? NNA-Berichte 12, 24-30.

Gliemann, L. (1973). Die Grauammer: *Emberiza calandra*. Verlag A. Ziemsen.

Hegelbach, J.F. (1984). Untersuchungen an einer Population der Grauammer (*Emberiza calandra* L.): Territorialität, Brutbiologie, Paarbindungssystem, Populationsdynamik und Gesangsdialekt. ADAG Administration & Druck AG.

Hölker, M. and Klähr, S. (2004). Bestandsentwicklung, Bruterfolg, Habitat und Nestlingsnahrung der Grauammer Miliaria calandra in der ackerbaulich intensiv genutzten Feldlandschaft der Hellwegbörde, Nordrhein-Westfalen. Charadrius 40, 133-151.

Perkins, A.J., Maggs, H.E. and Wilson, J.D. (2015). Crop sward structure explains seasonal variation in nest site selection and informs agri-environment scheme design for a species of high conservation concern: the Corn Bunting *Emberiza calandra*. Bird Study 62, 474-485.

Ryves, L. and Ryves, B. (1934). The breeding-habits of the Corn-bunting as observed in North Cornwall: with special reference to its polygamous habit. Brit. Birds 28, 2-26.

Sacher, T. and Bauschmann, G. (2011). Artenhilfskonzept für die Grauammer (*Miliaria calandra*) in Hessen. – Gutachten im Auftrag der Staatlichen Vogelschutzwarte für Hessen, Rheinland-Pfalz und das Saarland.

Setchfield, R.P., Mucklow, C., Davey, A., Bradter, U. and Anderson, G.Q. (2012). An agri-environment option boosts productivity of Corn Buntings *Emberiza calandra* in the UK. Ibis 154, 235-247.

Stein-Bachinger, K., Fuchs, S. and Gottwald, F. (2010). Naturschutzfachliche Optimierung des ökologischen Landbaus" Naturschutzhof Brodowin": Ergebnisse des E+ E-Projektes" Naturschutzhof Brodowin". – Bundesamt für Naturschutz, Bonn Bad Godesberg.

Suter, C., Rehsteiner, U. and Zbinden, N. (2002). Habitatwahl und Bruterfolg der Grauammer *Miliaria calandra* im Grossen Moos. Ornithologischer Beobachter 99, 105-115.

Preprtint - version July 2024

### Supplemental statistical analyses

#### Statistical supplement A – Results for Clutch size and Hatching rate models

The figures and tables report estimates from generalized linear models predicting variation in Corn Bunting clutch size (log-link based on a generalized Poisson distribution) and in hatching rates (logit-link based on a binomial distribution).

##### (A.1) Graphical results display

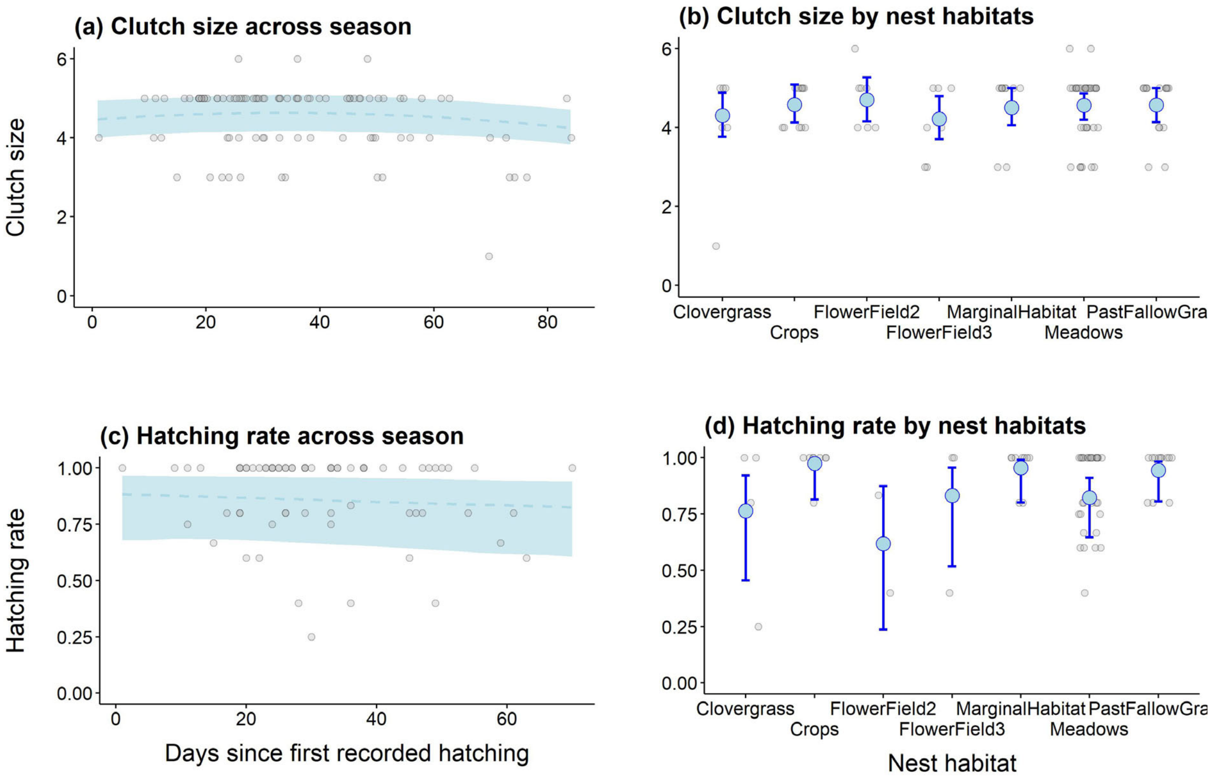

#### (A.2) Coefficient estimates from full models

<u>Model structures</u>:

```
ClutchSize ~ poly(HatchingDate, 2) + Nest habitat + Region,
            family = genpois(link = "log")

HatchingRate ~ poly(HatchingDate, 2) + ClutchSize +
            Nest habitat + Region,
            weights = ClutchSize,
            family = binomial(link = "logit")
```

*Table A.2a: Model parameter coefficient estimates.*

| Parameter | Coefficient* | SE | lwr.CI | upr.CI |
|---|---|---|---|---|
| ***(a) Clutch size model (log link)*** | | | | |
| (Intercept) [Nest habitat: alfalfa; Region: other] | 1.455 | 0.073 | 1.313 | 1.598 |
| Hatching day (linear) | -0.245 | 0.158 | -0.554 | 0.064 |
| Hatching day (quadratic) | -0.319 | 0.144 | -0.602 | -0.036 |
| Nest habitat: crops | 0.062 | 0.077 | -0.089 | 0.213 |
| Nest habitat: Flower fields (year 2) | 0.088 | 0.079 | -0.068 | 0.243 |
| Nest habitat: Flower fields (year 3+) | -0.021 | 0.090 | -0.197 | 0.156 |
| Nest habitat: seminatural margin habitat | 0.045 | 0.078 | -0.108 | 0.198 |
| Nest habitat: meadows | 0.058 | 0.063 | -0.066 | 0.182 |
| Nest habitat: Pasture, fallow grassland | 0.060 | 0.070 | -0.077 | 0.198 |
| Region: Rottenburg | -0.005 | 0.046 | -0.095 | 0.085 |
| ***(b) Hatching rate model (logit link)*** | | | | |
| (Intercept) [Nest habitat: alfalfa; Region: other] | 1.214 | 1.465 | -1.657 | 4.084 |
| Hatching day (linear) | -2.375 | 1.575 | -5.461 | 0.712 |
| Hatching day (quadratic) | 0.132 | 1.678 | -3.158 | 3.421 |
| Clutch size | -0.055 | 0.282 | -0.607 | 0.498 |
| Nest habitat: crops | 2.464 | 1.181 | 0.150 | 4.778 |
| Nest habitat: Flower fields (year 2) | -0.697 | 0.936 | -2.532 | 1.138 |
| Nest habitat: Flower fields (year 3+) | 0.425 | 0.944 | -1.425 | 2.275 |
| Nest habitat: seminatural margin habitat | 1.863 | 0.952 | -0.002 | 3.729 |
| Nest habitat: meadows | 0.357 | 0.635 | -0.888 | 1.602 |
| Nest habitat: Pasture, fallow grassland | 1.654 | 0.838 | 0.012 | 3.296 |
| Region: Rottenburg | 0.444 | 0.550 | -0.634 | 1.523 |

** Note: all coefficient estimates given as differences in the log ratio (model a) and in the log of odds (model b).*

Preprint - version July 2024

#### (A.3) Pairwise comparisons among nest habitats

| Contrasts | [odds]ratio* | SE | lower.CI | upper.CI |
|---|---|---|---|---|
| ***(a) Clutch size model (log link)*** | | | | |
| Alfalfa.Clover.grass - Crops | 0.940 | 0.072 | 0.808 | 1.093 |
| Alfalfa.Clover.grass - Flower fields (year 2) | 0.916 | 0.073 | 0.784 | 1.070 |
| Alfalfa.Clover.grass - Flower fields (year 3+) | 1.021 | 0.092 | 0.856 | 1.218 |
| Alfalfa.Clover.grass - Marginal.habitats | 0.956 | 0.075 | 0.820 | 1.114 |
| Alfalfa.Clover.grass - Meadow | 0.944 | 0.060 | 0.834 | 1.068 |
| Alfalfa.Clover.grass - Pasture.Fallowgrassland | 0.941 | 0.066 | 0.820 | 1.081 |
| Crops - Flower fields (year 2) | 0.975 | 0.072 | 0.843 | 1.127 |
| Crops - Flower fields (year 3+) | 1.087 | 0.082 | 0.938 | 1.259 |
| Crops - Marginal.habitats | 1.017 | 0.069 | 0.890 | 1.162 |
| Crops - Meadow | 1.005 | 0.058 | 0.898 | 1.124 |
| Crops - Pasture.Fallowgrassland | 1.002 | 0.062 | 0.888 | 1.130 |
| Flower fields (year 2) - Flower fields (year 3+) | 1.115 | 0.095 | 0.943 | 1.317 |
| Flower fields (year 2) - Marginal.habitats | 1.044 | 0.077 | 0.904 | 1.205 |
| Flowerfield.2 - Meadow | 1.031 | 0.060 | 0.919 | 1.156 |
| Flower fields (year 2) - Pasture.Fallowgrassland | 1.028 | 0.068 | 0.903 | 1.170 |
| Flower fields (year 3+) - Marginal.habitats | 0.936 | 0.071 | 0.806 | 1.087 |
| Flower fields (year 3+) - Meadow | 0.924 | 0.067 | 0.801 | 1.067 |
| Flower fields (year 3+) - PastureFallow.grassland | 0.922 | 0.072 | 0.792 | 1.074 |
| Marginal.habitats - Meadow | 0.987 | 0.056 | 0.884 | 1.103 |
| Marginal.habitats - Pasture.Fallowgrassland | 0.985 | 0.063 | 0.868 | 1.118 |
| Meadow - Pasture.Fallowgrassland | 0.997 | 0.046 | 0.911 | 1.092 |
| ***(b) Hatching rate model (logit link)*** | | | | |
| Alfalfa.Clover.grass - Crops | 0.085 | 0.100 | 0.008 | 0.861 |
| Alfalfa.Clover.grass - Flower fields (year 2) | 2.008 | 1.880 | 0.321 | 12.583 |
| Alfalfa.Clover.grass - Flower fields (year 3+) | 0.654 | 0.617 | 0.103 | 4.157 |
| Alfalfa.Clover.grass - Marginal.habitats | 0.155 | 0.148 | 0.024 | 1.002 |
| Alfalfa.Clover.grass - Meadow | 0.700 | 0.445 | 0.201 | 2.431 |
| Alfalfa.Clover.grass - Pasture.Fallowgrassland | 0.191 | 0.160 | 0.037 | 0.988 |
| Crops - Flower fields (year 2) | 23.598 | 30.210 | 1.919 | 290.132 |
| Crops - Flower fields (year 3+) | 7.681 | 9.594 | 0.664 | 88.838 |
| Crops - Marginal.habitats | 1.823 | 2.284 | 0.156 | 21.245 |
| Crops - Meadow | 8.222 | 8.888 | 0.988 | 68.417 |
| Crops - Pasture.Fallowgrassland | 2.247 | 2.720 | 0.209 | 24.104 |
| Flower fields (year 2) - Flower fields (year 3+) | 0.326 | 0.362 | 0.037 | 2.879 |
| Flower fields (year 2) - Marginal.habitats | 0.077 | 0.083 | 0.009 | 0.634 |
| Flowerfield.2 - Meadow | 0.348 | 0.257 | 0.082 | 1.476 |
| Flower fields (year 2) - Pasture.Fallowgrassland | 0.095 | 0.087 | 0.016 | 0.568 |
| Flower fields (year 3+) - Marginal.habitats | 0.237 | 0.241 | 0.032 | 1.737 |
| Flower fields (year 3+) - Meadow | 1.070 | 0.905 | 0.204 | 5.618 |
| Flower fields (year 3+) - PastureFallow.grassland | 0.293 | 0.286 | 0.043 | 1.991 |
| Marginal.habitats - Meadow | 4.511 | 3.731 | 0.892 | 22.820 |
| Marginal.habitats - Pasture.Fallowgrassland | 1.233 | 1.213 | 0.179 | 8.480 |
| Meadow - Pasture.Fallowgrassland | 0.273 | 0.177 | 0.077 | 0.975 |

** Note: all coefficient estimates are given as log ratios (model a) and odds ratios (model b).*

Preprint - version July 2024

#### Statistical supplement B – Results for first egg day (FED) models

The tables report coefficient estimates for Gaussian linear models predicting variation in Corn Bunting first egg days (FED) with region and nesting habitat.

##### (AB1) Overall differences in first egg day (FED) between study regions

Model structure:

```
FED ~ Region + (1|Year), family = gaussian
```

*Table B.1a: Model parameter coefficient estimates.*

| Parameter | Coefficient* | SE | lwr.CI | upr.CI |
|---|---|---|---|---|
| (Intercept) [Creglingen] | 158.083 | 3.497 | 151.230 | 164.937 |
| Ertingen.Klettgau | -1.715 | 5.525 | -12.544 | 9.114 |
| Rhine valley | -17.057 | 3.278 | -23.481 | -10.633 |
| Rottenburg | -12.108 | 3.117 | -18.217 | -6.000 |

** Note: all coefficient estimates are given as (differences in) day of the year (DOY).*

*Table B.1b: Posterior pairwise comparisons among regions.*

| Contrasts | difference* | lower.CI | upper.CI |
|---|---|---|---|
| Creglingen - Ertingen.Klettgau | 1.715 | -9.186 | 12.617 |
| Creglingen - Rhine valley | 17.057 | 10.589 | 23.524 |
| Creglingen - Rottenburg | 12.108 | 5.959 | 18.258 |
| Ertingen.Klettgau - Rhine valley | 15.341 | 4.953 | 25.729 |
| Ertingen.Klettgau - Rottenburg | 10.393 | 0.006 | 20.779 |
| Rhine valley - Rottenburg | -4.949 | -10.477 | 0.580 |

* Note: all contrasts are given as difference in day of the year (DOY).

Preprint - version July 2024

#### (B.2) Within-region models: differences in FED between nesting habitats

Model structure:

```
FED ~ Habitat + (1|Year), family = gaussian
```

*Table B.2a: Model parameter coefficient estimates.*

| Parameter | Coefficient* | SE | lwr.CI | upr.CI |
|---|---|---|---|---|
| ***(a) Within Region = Rottenburg*** | | | | |
| (Intercept) [Alfalfa & Clover grass] | 145.992 | 4.217 | 137.727 | 154.258 |
| Cereals | 4.028 | 5.484 | -6.720 | 14.777 |
| Flower fields | 14.104 | 6.172 | 2.008 | 26.200 |
| Marginal habitats | 6.019 | 6.679 | -7.072 | 19.110 |
| Meadows | -5.167 | 4.364 | -13.721 | 3.387 |
| Pasture | -2.202 | 5.269 | -12.530 | 8.126 |
| ***(b) Within Region = Rhine valley*** | | | | |
| (Intercept) [Fallow grassland] | 142.474 | 3.075 | 136.447 | 148.500 |
| Marginal habitats | -3.585 | 5.423 | -14.214 | 7.045 |
| Meadows | -4.402 | 4.721 | -13.655 | 4.850 |

** Note: all coefficient estimates are given as (differences in) day of the year (DOY).*

*Table B.2b: Posterior pairwise comparisons among nest habitats.*

| Contrasts | difference* | lower.CI | upper.CI |
|---|---|---|---|
| ***(a) Contrasts within Rottenburg region*** | | | |
| Alfalfa.Clover-grass - Cereals | -4.028 | -14.914 | 6.857 |
| Alfalfa.Clover-grass - Flower.field | -14.104 | -26.355 | -1.853 |
| Alfalfa.Clover-grass - Marginal.habitats | -6.019 | -19.277 | 7.239 |
| Alfalfa.Clover-grass - Meadow | 5.167 | -3.496 | 13.830 |
| Alfalfa.Clover-grass - Pasture | 2.202 | -8.257 | 12.662 |
| Cereals - Flower.field | -10.076 | -22.948 | 2.797 |
| Cereals - Marginal.habitats | -1.991 | -15.766 | 11.785 |
| Cereals - Meadow | 9.195 | 0.237 | 18.154 |
| Cereals - Pasture | 6.231 | -4.036 | 16.497 |
| Flower.field - Marginal.habitats | 8.085 | -6.194 | 22.364 |
| Flower.field - Meadow | 19.271 | 8.982 | 29.560 |
| Flower.field - Pasture | 16.306 | 4.398 | 28.215 |
| Marginal.habitats - Meadow | 11.186 | -0.440 | 22.812 |
| Marginal.habitats - Pasture | 8.221 | -4.859 | 21.302 |
| Meadow - Pasture | -2.965 | -10.889 | 4.960 |
| ***(b) Contrasts within Rhine valley region*** | | | |
| Fallow.grassland - Marginal.habitats | 3.585 | -7.404 | 14.574 |
| Fallow.grassland - Meadow | 4.402 | -5.163 | 13.967 |
| Marginal.habitats - Meadow | 0.817 | -10.785 | 12.420 |

* Note: all contrasts are given as difference in day of the year (DOY).

Preprint - version July 2024

#### Statistical supplement C – Results for apparent nest survival models

Tables report coefficient estimates for linear models with binomial error families and logit-link to predict variation in apparent survival between nests depending on nest habitat, nest visitation, and agricultural land management.

##### (C.1) Variation in apparent survival between nest habitats

Model structure:

```
Nest.success ~ Habitat + (1|Year), family = binomial(link = "logit")
```

*Table C.1a: Model parameter coefficient estimates.*

| Parameter | Coefficient* | SE | lwr.CI | upr.CI |
|---|---|---|---|---|
| (Intercept) [Alfalfa & Clover grass] | -0.369 | 0.444 | -1.239 | 0.501 |
| Cereals | 0.864 | 0.548 | -0.209 | 1.938 |
| Fallow grassland | 1.518 | 0.792 | -0.034 | 3.070 |
| Flower fields (year 2) | -0.849 | 0.922 | -2.656 | 0.957 |
| Flower fields (year 3+) | 1.050 | 0.836 | -0.590 | 2.689 |
| Marginal habitats | 0.592 | 0.647 | -0.675 | 1.859 |
| Meadows | 0.975 | 0.507 | -0.020 | 1.969 |
| Pasture | 0.984 | 0.641 | -0.274 | 2.241 |
| Rape and root crops | 1.464 | 1.240 | -0.966 | 3.895 |

** Note: coefficient estimates in this logit-link model are given as (differences in) the log of odds.*

*Table C.1b Posterior contrasts in apparent nest survival among nest habitats.*

| Contrasts | odds ratio* | lower.CI | upper.CI |
|---|---|---|---|
| Alfalfa.Clover.grass - Cereals | 0.421 | 0.144 | 1.233 |
| Alfalfa.Clover.grass - Fallow.grassland | 0.219 | 0.046 | 1.034 |
| Alfalfa.Clover.grass - Flower fields (year 2) | 2.338 | 0.384 | 14.235 |
| Alfalfa.Clover.grass - Flower fields (year 3+) | 0.350 | 0.068 | 1.803 |
| Alfalfa.Clover.grass - Marginal.habitats | 0.553 | 0.156 | 1.964 |
| Alfalfa.Clover.grass - Meadow | 0.377 | 0.140 | 1.020 |
| Alfalfa.Clover.grass - Pasture | 0.374 | 0.106 | 1.315 |
| Alfalfa.Clover.grass - Rape.Rootcrops | 0.231 | 0.020 | 2.628 |
| Cereals - Fallow.grassland | 0.520 | 0.118 | 2.295 |
| Cereals - Flower fields (year 2) | 5.548 | 0.968 | 31.795 |
| Cereals - Flower fields (year 3+) | 0.831 | 0.173 | 3.996 |
| Cereals - Marginal.habitats | 1.313 | 0.407 | 4.238 |
| Cereals - Meadow | 0.896 | 0.371 | 2.160 |
| Cereals - Pasture | 0.887 | 0.273 | 2.889 |
| Cereals - Rape.Rootcrops | 0.549 | 0.050 | 6.062 |
| Fallow grassland - Flower fields (year 2) | 10.667 | 1.338 | 85.030 |
| Fallow grassland - Flower fields (year 3+) | 1.597 | 0.234 | 10.919 |
| Fallow grassland - Marginal.habitats | 2.524 | 0.509 | 12.510 |
| Fallow grassland - Meadow | 1.722 | 0.410 | 7.235 |
| Fallow grassland - Pasture | 1.706 | 0.326 | 8.918 |

Preprint – version July 2024

| | | | |
|---|---|---|---|
| Fallow grassland - Rape.Rootcrops | 1.055 | 0.076 | 14.686 |
| Flower fields (year 2) - Flower fields (year 3+) | 0.150 | 0.018 | 1.270 |
| Flower fields (year 2) - Marginal.habitats | 0.237 | 0.037 | 1.528 |
| Flowerfield.2 - Meadow | 0.161 | 0.030 | 0.870 |
| Flower fields (year 2) - Pasture | 0.160 | 0.025 | 1.018 |
| Flower fields (year 2) - Rape.Rootcrops | 0.099 | 0.006 | 1.644 |
| Flower fields (year 3+) - Marginal.habitats | 1.581 | 0.290 | 8.623 |
| Flower fields (year 3+) - Meadow | 1.078 | 0.236 | 4.925 |
| Flower fields (year 3+) - Pasture | 1.068 | 0.191 | 5.975 |
| Flower fields (year 3+) - Rape.Rootcrops | 0.661 | 0.045 | 9.798 |
| Marginal.habitats - Meadow | 0.682 | 0.225 | 2.066 |
| Marginal.habitats - Pasture | 0.676 | 0.169 | 2.701 |
| Marginal.habitats - RapeRootcrops | 0.418 | 0.035 | 4.980 |
| Meadow - Pasture | 0.991 | 0.344 | 2.852 |
| Meadow - Rape.Rootcrops | 0.613 | 0.057 | 6.566 |
| Pasture - Rape.Rootcrops | 0.618 | 0.051 | 7.571 |

* Note: all contrasts are given as survival odds ratios.

Preprint - Version July 2024

#### (C.2) Variation in predicted survival of land use activity between nest habitats

Model structure:

```
Landuse.survival ~ Habitat + (1|Year), weights = N_cases,
         family = betabinomial(link = "logit")
```

*Table C.2a: Model parameter coefficient estimates.*

| Parameter | Coefficient* | SE | lwr.CI | upr.CI |
|---|---|---|---|---|
| (Intercept) [Alfalfa & Clover grass] | -0.576 | 0.280 | -1.124 | -0.028 |
| Meadows | 0.331 | 0.323 | -0.301 | 0.963 |
| Pasture | 1.085 | 0.405 | 0.291 | 1.880 |

** Note: coefficient estimates in this logit-link model are given as (differences in) the log of odds.*

*Table C.2b Posterior contrasts in predicted survival of land use activity among nest habitats.*

| Contrasts | odds ratio* | lower.CI | upper.CI |
|---|---|---|---|
| (Alfalfa.Clover-grass) / Meadow | 0.718 | 0.382 | 1.352 |
| (Alfalfa.Clover-grass) / Pasture | 0.338 | 0.153 | 0.748 |
| Meadow / Pasture | 0.470 | 0.244 | 0.906 |

* Note: all contrasts are given as survival odds ratios.

#### (C.3) Variation in apparent survival of visited and non-visited nests

Model structure:

```
Nest.success ~ Visited + (1|Year), family = binomial(link = "logit")
```

*Table C.3a: Model parameter coefficient estimates.*

| Parameter | Coefficient* | SE | lwr.CI | upr.CI |
|---|---|---|---|---|
| (Intercept) [Nest not visited] | 0.074 | 0.256 | -0.428 | 0.576 |
| Nest visited at least once | 0.729 | 0.323 | 0.095 | 1.363 |

** Note: coefficient estimates in this logit-link model are given as (differences in) the log of odds.*

*Table C.3b Posterior contrasts in predicted survival depending on nest visitation.*

| Contrasts | odds ratio* | lower.CI | upper.CI |
|---|---|---|---|
| not visited / visited at least once | 0.482 | 0.256 | 0.909 |

* Note: contrasts are given as survival odds ratios.

Preprtint - version July 2024

### Raw data supplement

#### Documentation of raw data table columns

A raw data file (*.xlsx, *.csv) is provided separately. The following table provides information on each data column in the raw data file.

| Column name | Content |
|---|---|
| Year | Year of data collection |
| Region | Study region (4 levels, as given in supplemental Table S1) |
| Region_Mayfield | Study region (2 levels, simplified as used for Mayfield DSR analysis) |
| FirstEggDate | FED: Absolute date of first egg laying (observed or backcalculated as given in manuscript) |
| FirstEggDay_DOY | FED: expressed as Day of the Year (DOY) |
| HatchDate | Absolute data of hatching (observed or backcalculated as given in manuscript. Used in analysis of Mayfield DSR) |
| NestHabitat | Habitat or land use at nesting site. |
| FirstBrood | Categorization as first brood (1) or any later (replacement or second) broods (0). Inferred as plausible as possible based on previous obserations in same territory. |
| NestSuccess | TRUE: nest successful as inferred from parents feeding jumplings in vicinity of previously known nest. FALSE: nest failed, either directly confirmed or deduced from lack of jumpling feeding after expected date of nest leaving |
| Nest.visited | TRUE: nest visited at least once for confirmation, control of nest status. FALSE: All nest observations from distance, no neset visit |
| LandUseSurvival | Fraction of years in which this particular nest's breeding phase did not overlap with any land use activity (harvest, mowing, grazing) on the very same patch as documented in other years. |
| N_cases | Land use survival: number of years for which information on land use dates were available for this particular patch |
| Eggs.full | Clutch size (number of eggs) |
| Eggs.lost | Number of eggs lost from known clutch prior to hatching |
| Eggs.hatched | Number of eggs that hatched |
| Eggs.deaf | Number of unhatched eggs that remained in nest (infertile or non-developing for other reasons) |
| Clutch.Complete | Identifies nests with information on full clutch size |
| LastPres | Mayfield DSR: Day number (counted from earliest record in dataset) on which this nest was last present |
| LastCh | Mayfield DSR: Day number (counted from earliest record in dataset) on which this nest was last checked |
| VegHeight | Dominant vegetation height at nest site (in cm) |
| VegCover | Cover of herbaceous vegetation layer in 1m circle around the nest (in %) |
| NestHeight | Height of upper nest rim above ground (cm) |
| Analysis1.NestHabitat | Data included into the analysis of nest habitats |
| Analysis2.ClutchSize | Data included into the analysis of clutch sizes |
| Analysis3.HatchRate | Data included into the analysis of hatch rates |
| Analysis4.FED | Data included into the analysis of First Egg Dates (FED) |
| Analysis5.ApparentSurvival | Data included into the analysis of apparent survival |
| Analysis6.LandUseSurvival | Data included into the analysis of land use survival |
| Analysis7.DSR | Data included into the analysis of Daily nest Survival Rates (DSR) |

Preprint – version July 2024
